# Chronic Intermittent Ethanol and Lipopolysaccharide Exposure Differentially Alter Iba-1-Derived Microglia Morphology in the Prelimbic Cortex and Nucleus Accumbens Core

**DOI:** 10.1101/566034

**Authors:** BM Siemsen, JD Landin, JA McFaddin, KN Hooker, LJ Chandler, MD Scofield

## Abstract

Accumulating evidence has linked pathological changes associated with chronic alcohol exposure to neuroimmune signaling mediated by microglia. Prior characterization of the microglial structure-function relationship demonstrates that alterations in activity states occur concomitantly with reorganization of cellular architecture. Accordingly, gaining a better understanding of microglial morphological changes associated with ethanol exposure will provide valuable insight into how neuroimmune signaling may contribute to ethanol-induced reshaping of neuronal function. Here we have used Iba-1-staining combined with high-resolution confocal imaging and 3D reconstruction to examine microglial structure in the prelimbic (PL) cortex and nucleus accumbens (NAc) in male Long-Evans rats. Rats were either sacrificed at peak withdrawal following 14 days of exposure to chronic intermittent ethanol (CIE) or 48 hours after exposure to the immune activator lipopolysaccharide (LPS). LPS exposure resulted in dramatic structural reorganization of microglia in the PL cortex; including increased soma volume, overall cellular volume, and branching complexity. In comparison, CIE exposure was associated with a subtle increase in somatic volume and differential effects on microglia processes, which were largely absent in the NAc. These data reveal that microglial activation following a neuroimmune challenge with LPS or exposure to chronic alcohol exhibit distinct morphometric profiles and brain-region dependent specificity.

## Introduction

Microglia participate in neuroimmune signaling induced by exposure to environmental insults by releasing cytokines and neuromodulators to support healthy central nervous system function and to regulate synaptic transmission [1]. However, dysfunctional cognitive processes that occur during aging, neurodegenerative disorders, and alcohol use disorder (AUD), have been linked to aberrant or dysfunctional activation of microglia in specific brain regions [2, 3]. AUDs in particular represent a significant public health concern that places an immense burden on both individuals and society [4]. Rodent models of chronic alcohol exposure, which reproduce the somatic symptoms of physical dependence and withdrawal observed in humans [5], also engage neuropathological processes that parallel what has been observed in brains from human alcoholics. Apart from the established link between alcohol-induced neuroplastic changes in neurotransmitter systems such as glutamate, GABA, and dopamine (for review see [6]), there is accumulating clinical [7–9] and preclinical [10, 11] evidence for the role of altered neuroimmune activity in the neuroadaptations induced by chronic ethanol exposure that may facilate dysfunctional, frontal cortical-dependent, cognitive processes. As an example, immunoreactivity of ionized calcium binding adaptor molecule 1 (Iba-1), a protein specifically expressed in microglia [12], is increased in post-mortem brains of alcoholic patients [13].

Microglia express several membrane receptors that respond to molecular patterns associated with damage or pathogens, leading to an innate or adaptive immune response [14]. The superfamily of toll-like receptors (TLRs) are particularly important for recognizing extracellular signals linked to cellular damage and promoting the release of cytokines in response [15]. Canonically, systemic injections of lipopolysaccharide (LPS) have been utilized to increase TLR4-dependent central microglia activation [16]. Interestingly, several psychoactive compounds, such as opioids and cocaine, also activate microglia by binding to and activating TLR4, leading to central immune signaling [17, 18]. Akin to opioids and cocaine, studies have shown that ethanol exposure leads to microglia activation by increasing TLR4 and tumor necrosis factor alpha (TNFα) receptor-dependent neuroimmune signaling [19, 20]. Interestingly, ethanol exposure during adolescence increases TLR4 expression and increases the abundance of ligands for TLR4, including the damage-associated high-mobility group box (HMGB1) [21]. However, the degree to which ethanol leads to TLR4-dependent neurotoxic insult is currently under debate [11, 22–24]. Nevertheless, knock out studies support a TLR4-based mechanism for ethanol’s action on microglia, as loss of TLR4 prevents ethanol-induced microglial upregulation in the cortex and cytokine release in culture [25]. Apart from TLR4, ethanol also acts directly on NADPH-oxidase [26], inducing the release of reactive oxygen species from microglia, which have been linked to alcohol-induced neurodegeneration [27]. Further, systemic administration of microglia activation inhibitor minocyline has been shown to decrease ethanol self-administration [28]. Collectively, these data support a role for microglia activation in the ethanol-induced neuroimmune and neuroinflammatory cascades linked to the disruption of cognitive processes mediating the maintenance of escalated ethanol drinking.

As Iba-1 is specifically expressed in microglia in the CNS [12, 29], traditional immunohistochemical labeling and detection of Iba-1 has provided valuable insight into brain region-specific adaptations in microglia structure and proliferation following ethanol exposure. Several studies have shown that chronic ethanol exposure in rats increases the number of Iba-1 positive cells, or increases Iba-1 signal intensity, in a region-specific manner. For example, while six months of voluntary consumption of a 20% aqueous ethanol solution followed by 2 months of withdrawal did not alter the total number of microglia in the hippocampus, it did increase the proportion of activated microglia as indicated by elevated CD11b immunoreactivity [22]. Studies examining large numbers of microglia at relatively low magnification have reported that Iba-1 intensity is increased in a TLR4-dependent manner within a number of brain regions, including the PFC. This was observed following 35 days of chronic intermittent ethanol exposure (CIE), which was diminished after 28 days of withdrawal [30]. As such, it is likely that ethanol-induced activation of microglia plays a role in the neuroadaptations associated with chronic alcohol exposure.

Microglia are exceptionally morphologically dynamic [31], with activation-associated alterations in soma size and branching patterns serving as structural biomarkers for neuroimmune activation [32–34]. Further, recent studies demonstrate brain region-specific heterogeneity in microglial morphology [35] and gene expression profiles [36], establishing brain region heterogeneity and supporting a clear link between microglia structure and function. Interestingly, even *within* brain regions individual microglia occupy a broad spectrum of activation states, and thus structural signatures [37]. Historically, microglia morphology has been used to clasify microglia into either an activated or a resting state. However, a binary two-state categorization is may be overly simplistic given that microglia constantly survey and respond to the surrounding microenvironment and likely respond by progressing through a continuum of differential activation states [38]. Microglial processes are uniquely plastic and traverse the parenchyma at a rapid rate of 1-3 µm per minute [38], influencing synaptic transmission as they sample the interstitial fluid and respond to stimuli with the release of neuromodulators [39].

The morphological signature of microglial cells tightly correlates with their functional states [40–42]. For instance, ramified microglia are characterized by thin and elongated processes and are canonically considered to be in a “resting” state. Environmental challenges that induce neural inflammation can transform microglia into an “ameboid” or “reactive” state, often characterized by an enlarged soma and thick or condensed processes. Once activated, microglia upregulate specific receptors and express secretory analogues that contribute to the defense of the central nervous system to environmental insults [43, 44]. In general, microglia are remarkably sensitive to neural insult, and thus environmental challenges can produce rapid and extensive adaptations in microglial structure and function [45].

Although a significant body of work exists regarding the impacts of ethanol on microglia and neuroimmune signaling (for review see [46]), to date a systematic analysis of microglia morphological adaptations following CIE exposure has yet to be comprehensively conducted using high-resolution confocal microscopy and high fidelity digital reconstruction. Moreover, the effects of ethanol withdrawal have not been directly compared to canonical inducers of microglia activation, such as LPS. Here we show that CIE exposure of male rats produces brain region-dependent alterations in microglia morphometric features that display overlapping as well as dichotomous alterations induced by sub-chronic (2-day) immune activation following systemic LPS administration.

## Materials and Methods

### Animal Subjects

Male Long-Evans rats (*N*=15) were obtained from Envigo (Indianapolis, IN). Upon arrival, rats were single housed in standard polycarbonate cages in a temperature-controlled vivarium and maintained on a 12 hr/12 hr reverse light/dark cycle (lights on at 2100 hours) with *ad libitum* access to food and water throughout the experiment. Male *Cx3cr1*^*CreER*^-EYFP transgenic mice (N=3) were purchased from Jackson Laboratories (#021160). These mice constitutively express EYFP in lieu of endogenous Cx3cr1 brain-wide in microglia [47]. All experiments were performed in accordance with guidelines for animal care established by National Institutes of Health and approved by MUSC’s Institutional Animal Care and Use Committee.

### 2.2 Chronic intermittent ethanol exposure

Chronic intermittent ethanol (CIE) exposure was initiated using a well-characterized vapor inhalation model [48]. In brief, adult male Long-Evans rats at post-natal day (PD) 90 were subjected to 15 consecutive days of intermittent ethanol vapor exposure. Each exposure day consisted of placing rats in standard housing cages into clear acrylic ethanol vapor exposure chambers (Plas Labs; Lansing, MI) for a period of 14 hrs/day. During the 10 hrs out of the chambers, the rats were returned to the vivarium. Rats were placed into the chambers at 1800 hours and removed at 0800 hours. Immediately upon removal from the vapor chambers, a 5-point behavioral intoxication scale was utilized to assess the level of intoxication [49]. Immediately prior to placing the rats back into the chambers following the 10 hr ethanol withdrawal period, the severity of withdrawal based upon somatic signs was assessed using a 5-point rating system [48, 50]. On days 1, 5, 10, and 15, tail blood (40 µl) was collected immediately upon removal from chambers for subsequent determination of blood ethanol concentration (BEC). Air-exposed control rats were treated similarly to the CIE-exposed rats except they were not placed into the ethanol vapor chambers.

### LPS injections

A subset of male Long-Evans rats received intraperitoneal injections of 1.0 mg/kg lipopolysaccharide (LPS) from Escherichia coli (Sigma-Aldrich, L2630) in sterile saline (0.9%) once per day for two consecutive days. Rats were then perfused 24 hours after the second injection of LPS.

### Immunohistochemistry

Mice or rats were heavily anesthetized with urethane (30% w/v) and then transcardially perfused with 120 ml (rats) or 20 ml (mice) of 0.1M PBS (pH 7.4) followed by 180 ml (rats) or 20 ml (mice) of fresh 4% paraformaldehyde (PFA, pH 7.4) (Fisher Scientific, Hampton NH). Brains were rapidly removed, post-fixed in 4% PFA for 24 hours, and then transferred to 0.1M PBS containing 0.2% sodium azide (w/v) and stored at 4ºC until sectioning. One hundred µm thick coronal sections containing the PL cortex (Rat coordinates: AP 2.76mm – AP 4.2mm from bregma, Mouse coordinates: AP 1.5mm – 2mm from bregma) and the NAc (Rat coordinates: AP 1.8mm – AP 0.7mm from bregma, Mouse coordinates: AP 1mm-1.4mm from bregma) were sliced on a vibrating microtome (Leica Biosystems, VT1000 S). Serial sections were stored in 0.1M PBS with 0.2% sodium azide until processing. Three to four sections containing the PL cortex or the NAc were processed for immunohistochemical fluorescent detection of Iba-1 or multiplex detection of Iba-1 and GFP. Sections were blocked in 0.1M PBS containing 2% Triton X-100 (PBST) and 2% normal goat serum (NGS, Jackson Immunoresearch, # 005-000-121) for 2 hours at room temperature with agitation. This was followed by incubation in PBST containing 2% NGS and rabbit anti-Iba-1 (1:1000, Wako #019-19741, RRID:AB_8939504) and chicken anti-GFP (1:1000, Abcam, Cambridge UK, #ab13970, RRID:AB_300798) primary antisera overnight at 4ºC with agitation. Sections were then washed 3 × 10 minutes in PBST, incubated in PBST containing 2% NGS with goat anti-rabbit secondary antisera conjugated to Alexa Fluor® 488 (1:1000, Abcam, Cambridge UK, #ab150169) and Alexa Fluor® 647 (1:1000, Abcam, Cambridge UK, #ab150079, RRID: AB_2722623) for five hours at room temperature, washed 3 × 10 minutes in PBST, and then mounted on superfrost-plus slides with ProLong™ Gold Antifade (Invitrogen, Carlsbad CA, #P36930). Mounted slides were stored at 4ºC until confocal imaging.

### High-resolution confocal microscopy

All imaging and analyses were performed by an investigator blind to experimental groups. Confocal Z-series image data sets of Iba-1+ microglia were captured using a Leica SP8 laser-scanning confocal microscope with a 63X (N.A. 1.4) oil-immersion objective. The parameters for imaging acquisition involved two separate modalities. The first set of parameters was designed to image networks of microglia for high throughput analyses of the somatic dimensions. These high throughput data sets were acquired using a diode 638 nm laser line, 2048 × 2048 frame size, 0.75X digital zoom, and a Z-step size of 0.37 µm. The entirety of all visible Iba-1 signal was collected throughout the thickness of the slice (data sets spanning 70-100 µm). The second set of imaging parameters, which were optimized for examining individual microglial cells at high resolution, involved image acquisition using the same objective and laser line, 2336 × 2336 frame size, 1.5X digital zoom, and a Z-step size of 0.24 µm. Data sets collected with this modality typically spanned 50-70 µm. For the high-resolution single-cell imaging approach, care was taken to ensure that the entirety of the cell was collected in the image by positioning the soma in the middle of the Z-stack during imaging. This ensured that all microglial processes for the cell of interest, projecting radially and axially, were completely collected. Microglia in the PL were imaged in layer V while those in the NAc were imaged in the core (NAcore) region dorsomedial to the anterior commissure.

### Imaris 3D reconstruction and analysis

Following confocal image acquisition, Z-series data sets were imported into Imaris software (Version 9.0, Bitplane, Belfast UK). For high throughput analyses of somatic dimensions, a smooth 3D space-filling model was constructed in order to best fit microglial cell bodies in a semi-automated manner. Inaccurate or falsely recognized somas were adjusted or removed, respectively, by an investigator blind to experimental groups. The total number of cells contained within a field, as well as each microglial soma volume, were exported.

Confocal data sets containing high-resolution Z-stacks of individual microglia were imported to Imaris for 3D modeling, skeletonization, branching, and Sholl analyses. Once an individual microglial cell with its reconstructed fine processes was isolated, the Filament module in Imaris was used to skeletonize each microglia in a semi-automated process. For this procedure, the thinnest process diameter was set to 0.5 µm and the overall seed size diameter was set to 8 µm. Each microglia skeletonization was then visually inspected and manually adjusted to best fit the Iba-1 signal with care. Next, the filament module was combined with a MATLAB plugin to produce a 3D convex hull for each cell. This shape is mathematically defined as the minimum polyhedron that can contain the filament resulting from skeletonization. This convex hull was used as an index of the 3D territory that each microglia occupied. Skeletonization analyses of the microglial processes included determination of 3D Sholl intersections obtained as a function of distance from the cells center (the middle of the soma), measurement of the convex hull volume (in µm^3^) as an index of territory occupied, the overall cellular volume of microglia (soma and processes), and determination of the number of filament branch points.

For comparison of Iba-1 immunohistochemical detection with the detection of immunoamplified EYFP in Cx3cr1 transgenic mice, 24 cells were imaged and isolated as described above. In this case, Iba-1 was used as the source channel for isolating individual cells, then a 3D space filling model was built on both Iba-1 and EYFP. The overall volume of each cell for each source channel was then compared. Three to four sections were imaged per animal.

### Statistical analyses

Statistical analyses were performed using GraphPad Prism software (version 6, GraphPad, San Diego CA). Normality of data was analyzed using Shapiro-Wilk test. If data within a group was found to be non-normally distributed, a corrected analysis was performed as specified in the results. BEC and withdrawal scores were analyzed with a repeated-measures one-way ANOVA followed by Dunnett’s multiple comparison test comparing days 10 and 15 to day 5. When three groups were compared across a single dependent variable, a one-way ANOVA was used. Tukey’s multiple comparison test was used when a significant effect of treatment (LPS or CIE versus control) was revealed. The number of Sholl intersections as a function of distance from soma start point was analyzed with a two-way repeated measures ANOVA with treatment (CIE or LPS versus Naïve) as a between-subjects variable and distance from soma center as a within-subjects variable. Bonferroni-corrected pairwise comparison test was used when a significant interaction was observed. For high-throughput analyses, the average soma volume was calculated for confocal data set collected, values were generated for 4-6 data sets per animal. These values were then used to generate an animal average, then animal averages were reported. Individual soma volume for each animal in each group were also plotted as independent data points to investigate population shifts in microglia soma volume as a function of treatment and analized with three individual Kolomogorov-Smirnov tests comparing control to CIE, control to LPS, and CIE to LPS. A paired student’s t-test was used when comparing the cell volume of Iba-1 compared to cell volume of Cx3cr1-EYFP

## Results

### Iba-1 labeling of microglia compared to transgenic mice

In the present study, we examined microglial morphology in the PL cortex and NAc of brain sections obtained from three experimental groups of rats: an ethanol-naïve control group, a group of rats that had been subjected to CIE and sacrificed 10 hours following the last ethanol exposure, and a group of rats that had been injected with LPS once daily for two consecutive days and then sacrificed 24 hours later (Figure 1).

**Figure 1.**
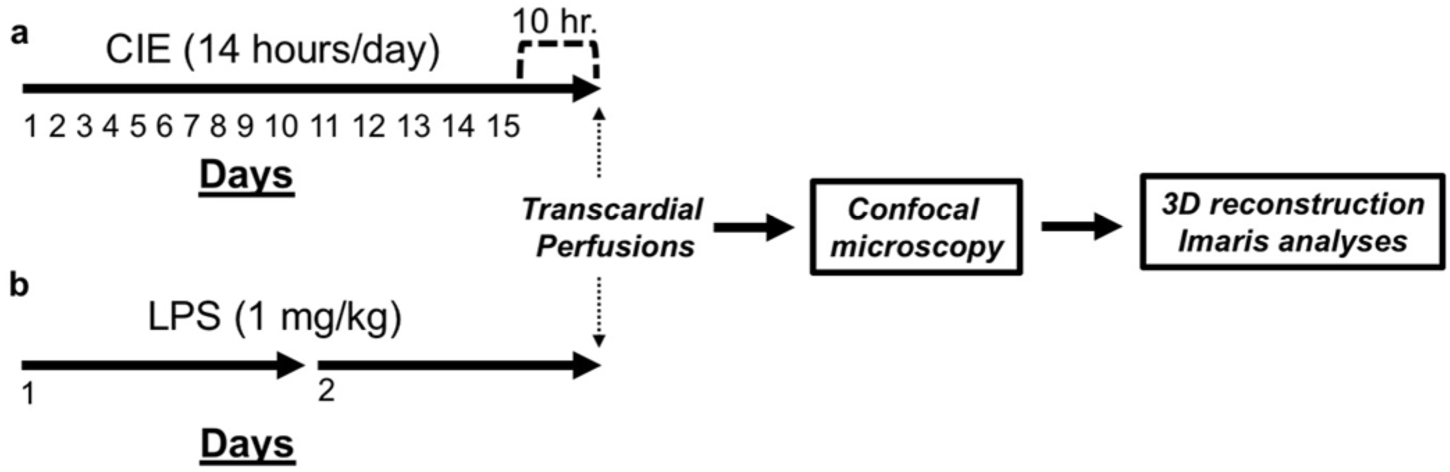
Schematic depiction of the experimental timeline. **a)** Timeline depicting the length of CIE and the sacrifice time-point during withdrawal. **b**) Timeline depicting the number of LPS injections received and the sacrifice time-point following LPS exposure. CIE or LPS treated rats were compared to ethanol-naïve controls on each measure.

We first performed a control experiment to demonstrate the fidelity of Iba-1 immunohistochemistry as a means to fully label microglia syncytia in both the PL cortex and the NAc. Figure 2a-b shows Iba-1 immunohistochemical detection (red) and EYFP (green) from Cx3cr1 transgenic mice in the PL cortex. 3-4 images of microglia were imaged per animal. Figure 2c-d shows the same immunohistochemical detection in the NAc and Figure 2e-f shows a higher magnification merge. As shown in Figure 2g, a two-tailed paired t-test indicated that the cell volume (in µm^3^) derived from the EYFP source channel (1708 +/− 143) was slightly lower than that of the Iba-1 source channel (1934 +/−202.1) in the PL cortex (t_(9)_=2.28, *p*=0.049). As shown in Figure 2h, the EYFP source channel (1466 +/−164) was also significantly lower than that of the Iba-1 source channel (1804+/−146.5) in the NAc (t_(7)_=4.67, *p*=0.002). Thus, Iba-1 immunohistochemistry produced a slightly more complete labeling of microglial cells than what was obtained following EYFP immunohistochemistry in both the PL and the NAcore of Cx3cr1 transgenic mice.

**Figure 2.**
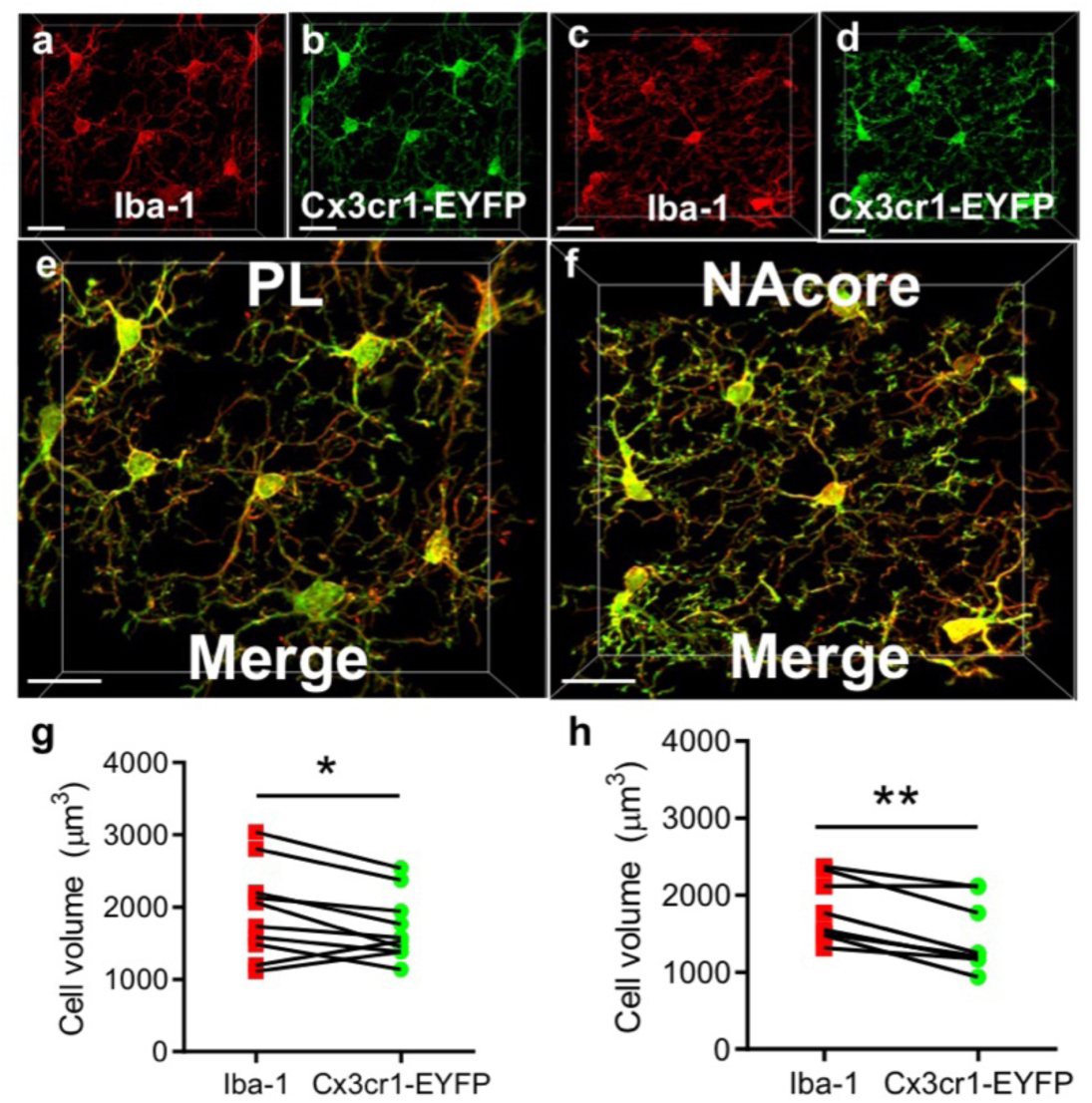
Cellular reconstructions of microglia from genetic and immunofluorescent labeling techniques produce similar results. **a-b)** Labeling produced by Iba-1 IHC (Red) and Cx3cr1-EYFP fluoresence (Green) in the PL portion of the PFC in Cx3cr1-EYFP mice (*n*=3). Isolation of microglia for cell volume analyses were done as described within the materials and methods section. **c-d)** Labeling produced by Iba-1 IHC and Cx3cr1-EYFP fluorescence in the NAcore in Cx3cr1-EYFP mice. **e-f)** High magnification merged image of data shown in **a-d**. **g-h)** Quantification of isolated cell volume using either Iba-1 or Cx3cr1-EYFP signal as the source channel in PL cortex (**g**) and NAcore (**h**). Scale bars=20 µm.

### Measurements of blood ethanol concentration, withdrawal scores, and representative immunohistochemical detection of Iba-1 in the PL cortex and NAcore

To obtain quantitative measures of alcohol intoxication and physical dependence, intoxication and withdrawal scoring were performed daily while BECs were obtained on 4 separate days of CIE exposure from tail-vein blood. As shown in Figure 3a, the BEC averaged across all 4 days was 294.4 +/− 19.67. The corresponding level of intoxication averaged across all days was 2.17 +/− 0.09 (Figure 3b), which equates to a mild-to-moderate level of intoxication based upon the parameters of the rating scale. A Geisser-Greenhouse-corrected repeated measures one-way ANOVA comparing withdrawal scores across time revealed a significant effect of time (F_(1.8,7.2)_=17.00, *p*=0.002, Figure 3c). Dunnett’s multiple comparison test indicated that rats on both day 10 (*p*=0.034) and day 15 (*p*=0.012) exhibited significantly higher withdrawal scores compared to day 5.

**Figure 3.**
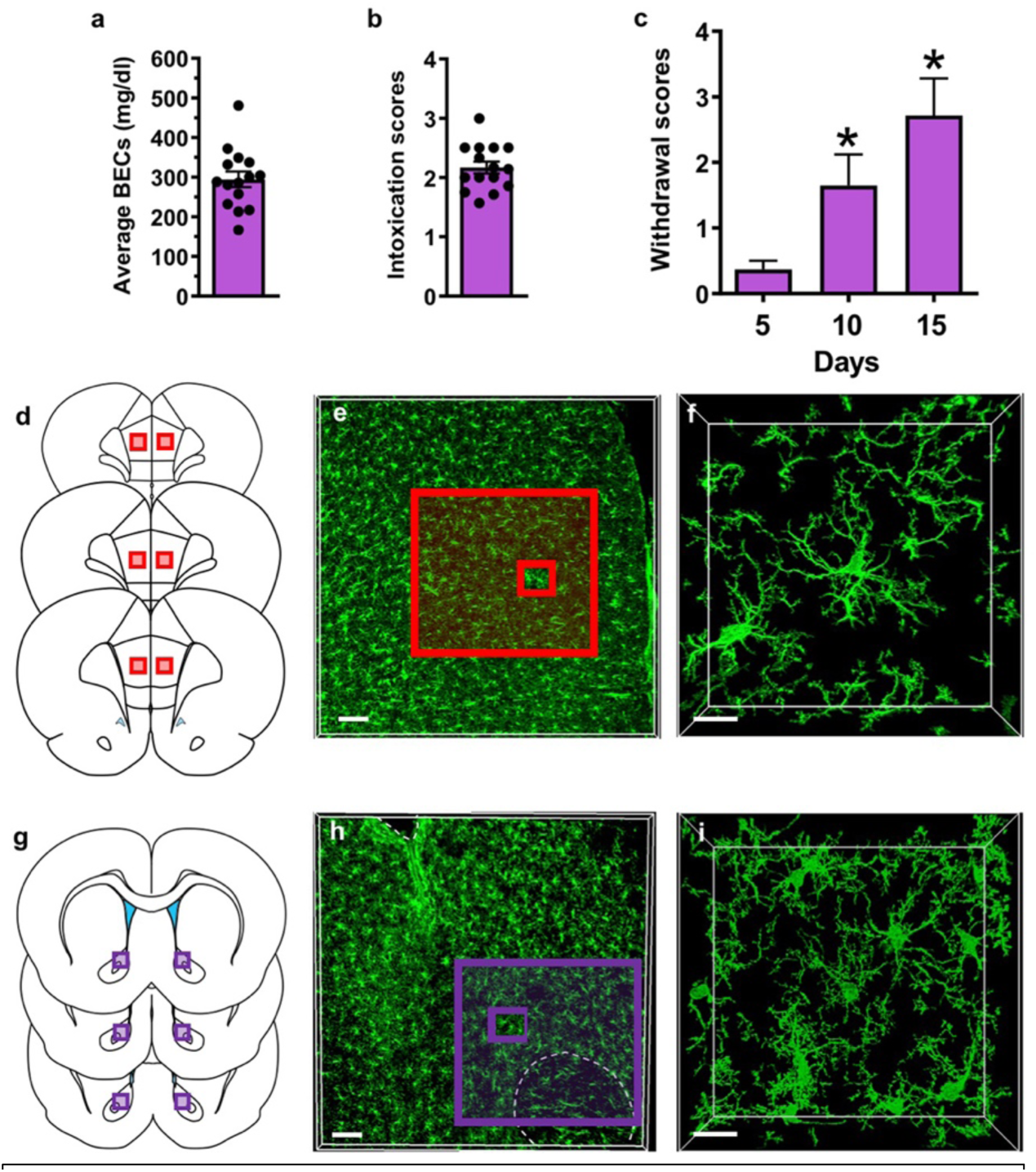
Assessment of ethanol exposure and representative Iba-1 signal in PL cortex and NAcore. **a**) Blood ethanol concentrations (BEC) and **b**) intoxication scores of CIE exposed rats averaged across day 5, 10, and 15 of the CIE exposure period. **c**) Somatic withdrawal scores assessed after 5, 10, and 15 days of CIE exposure (\**p*<0.05 compared to day 5). **d**) The PL subdivision of the medial prefrontal cortex was sampled along the anterior-posterior axis. **e**) Low magnification image of Iba-1 immunoreactivity in the PL cortex. **f**) High magnification image Iba-1 immunoreactive microglia in the PL cortex. The outer red boxed region in **e** represents the area that corresponds to the box region shown in **d**, while the smaller inset red box represents the image area shown in **f**. **g**) The NAcore subdivision of the nucleus accumbens was also sampled from sections across the anterior-posterior axis as described above. **e**) Low magnification images of Iba-1 immunoreactivity in the NAcore. **f**) Higher magnification image of Iba-1 immunoreactive microglia in the NAcore. As above, the outer purple boxed region in **h** represents the area that corresponds to the box region shown in **g** while the smaller inset purple box represents the image area shown in **i**. Scale bars=100 µm (e,h) and 15 µm (f,i).

Given that the PL cortex and NAcore are two key brain regions that undergo neuropathological changes in response to chronic alcohol exposure [51, 52], immunohistochemical labeling and subsequent confocal imaging of Iba-1+ microglia was carried out in sections containing these regions. Figure 3d shows the region sampled in the PL cortex and Figure 3e shows representative low magnification immunohistochemical detection of Iba-1 in the PL cortex whereas Figure 3f shows a representative higher magnification image of several Iba-1+ microglia. Figure 3g shows representative the region sampled in the NAcore and Figure 3h-i shows low and high magnification images of Iba-1 detection in the NAcore.

### High throughput, population-based analyses on microglia density and soma volume in the PL cortex and NAcore after CIE or LPS exposure

In the PL cortex, Iba-1 immunostaining produced robust labeling of microglia in sections obtained from control (Figure 4a, *n*=5), CIE-exposed (Figure 4b, *n*=5), and LPS injected rats (Figure 4c, *n*=5). Dimensions of the somatic compartment of microglia cells were analyzed using 3D space-filling models, which were color coded to indicate somatic volume in µm^3^. We imaged 4-6 fields of microglia in the PL cortex for each rat, and generated data for 5 rats for each condition, resulting in a total of 958 cells in the control group, 951 cells in CIE-exposed animals, and 902 cells in LPS-exposed animals. When these data were collapsed and expressed as an average across all rats in a group, a one-way ANOVA revealed a significant main effect of treatment on the number of cells/µm^3^ (F_(2,12)_=5.25, *p*=0.023). Tukey’s multiple comparison test indicated that CIE (*p*=0.024), but not LPS (*p*=0.791) exposure decreased the number of cells/µm^3^ when compared to control rats (Figure 4d). In addition, a significant main effect of treatment was also observed for somatic volume (F_(2,12)_=20.61, *p*=0.0001) (Figure 4e). Post-hoc analysis with Tukey’s multiple comparison tests indicated that while CIE did not significantly increase microglia soma volume relative to control rats (*p*=0.263), LPS treated rats exhibited a robust increase in soma volume relative to control (*p*=0.0001) and CIE-exposed rats (*p*=0.002). As shown in Figure S1A, cumulative frequency distributions of soma volume that presents each cell imaged and quantified as an individual data point, akin to the approach that was previously used to analyzed population shifts in dendritic spine morphology [53, 54]. Comparison of the microglial soma volume cumulative frequency distribution in the PL cortex of CIE exposed rats to the cumulative frequency distribution in control rats revealed a significant rightward shift in the somatic volume distribution (K-S *D*=0.13, *p*<0.0001). The fact that this difference was not present in the animal averages reflects the subtlety of CIE’s impact on soma volume. As expected, cumulative frequency distributions of LPS-exposed rats show a pronounced rightward shift in the soma volume distribution when compared to control animals (K-S *D*=0.31, *p*<0.0001) and to CIE-exposed rats (K-S *D*=0.20, *p*<0.0001). Immunostaining for Iba-1 in the NAcore also produced robust signal across all groups (Figure 5a-c). As with the PL, we imaged 4-6 fields of microglia for each animal, and imaged a total of 5 animals. This produced a total of 929 total cells in control rats, 833 total cells in CIE-exposed rats, and 942 total cells in LPS-exposed rats. Analysis of the immunostaining in the NAcore of the three experimental groups of rats yielded a significant main effect of treatment on the number of cells per µm^3^ when collapsed across animals (F_(2,12)_=6.72, *p*=0.011). Tukey’s multiple comparison test further indicated that LPS (*p*=0.009), but not CIE (*p*=0.295), exposure reduced the number of cells per µm^3^, when compared to control rats (Figure 5d), an effect that was opposite to that observed in the PL cortex. There was also a significant main effect of treatment on the average soma volume (F_(2,12)_=17.43, *p*=0.0003). Tukey’s multiple comparison test indicated that LPS exposure resulted in a significant increase in microglia soma volume when compared to control rats (*p*=0.0004) and CIE-exposed rats (*p*=0.001, Figure 5e). As described above for the PL cortex, each cell from each group was also expressed as an individual data point that resulted in a cumulative frequency distribution of soma volume (Figure S1b). As was the case with the PL cortex, comparison of CIE-exposed rats to control rats revealed a subtle rightward shift in the somatic volume distribution in the NAcore (K-S *D*=0.09, *p*<0.01). LPS exposure was again associated with a more dramatic rightward shift in the somatic volume frequency distribution of microglia in the NAcore compared to control (K-S *D*=0.32, *p*<0.0001) and CIE-exposed rats (K-S *D*=0.24, *p*<0.0001).

**Figure 4.**
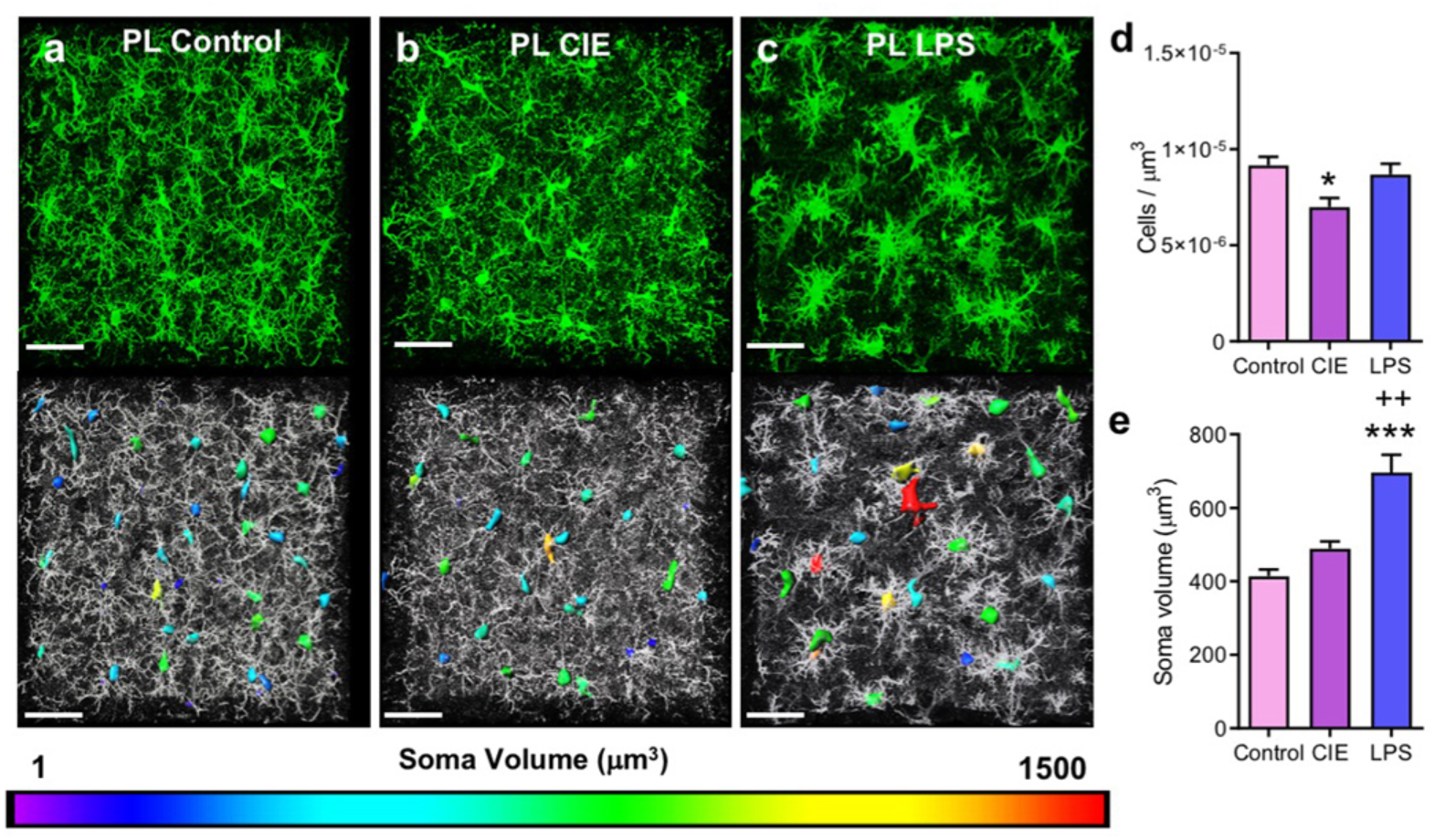
Both CIE and LPS exposure resulted in an increase in somatic volume of microglia in the PL cortex. Representative Iba-1 staining of microglia in the PL cortex for control (**a**), CIE-exposed (**b**), and LPS-exposed (**c**) rats. Somas are color-coded to indicate the spectrum of soma volumes observed for each treatment. **d**) CIE exposure reduced the number of Iba-1 immunoreactive microglia per micron^3^ in the PL cortex. **e**) LPS (but not CIE) exposure was associated with a significant increase in the somatic volume when compared to control rats. \**p*<0.05, \*\*\**p*<0.001 compared to control, ++*p*<0.01 compared to CIE. Scale bar=40 µm.

**Figure 5.**
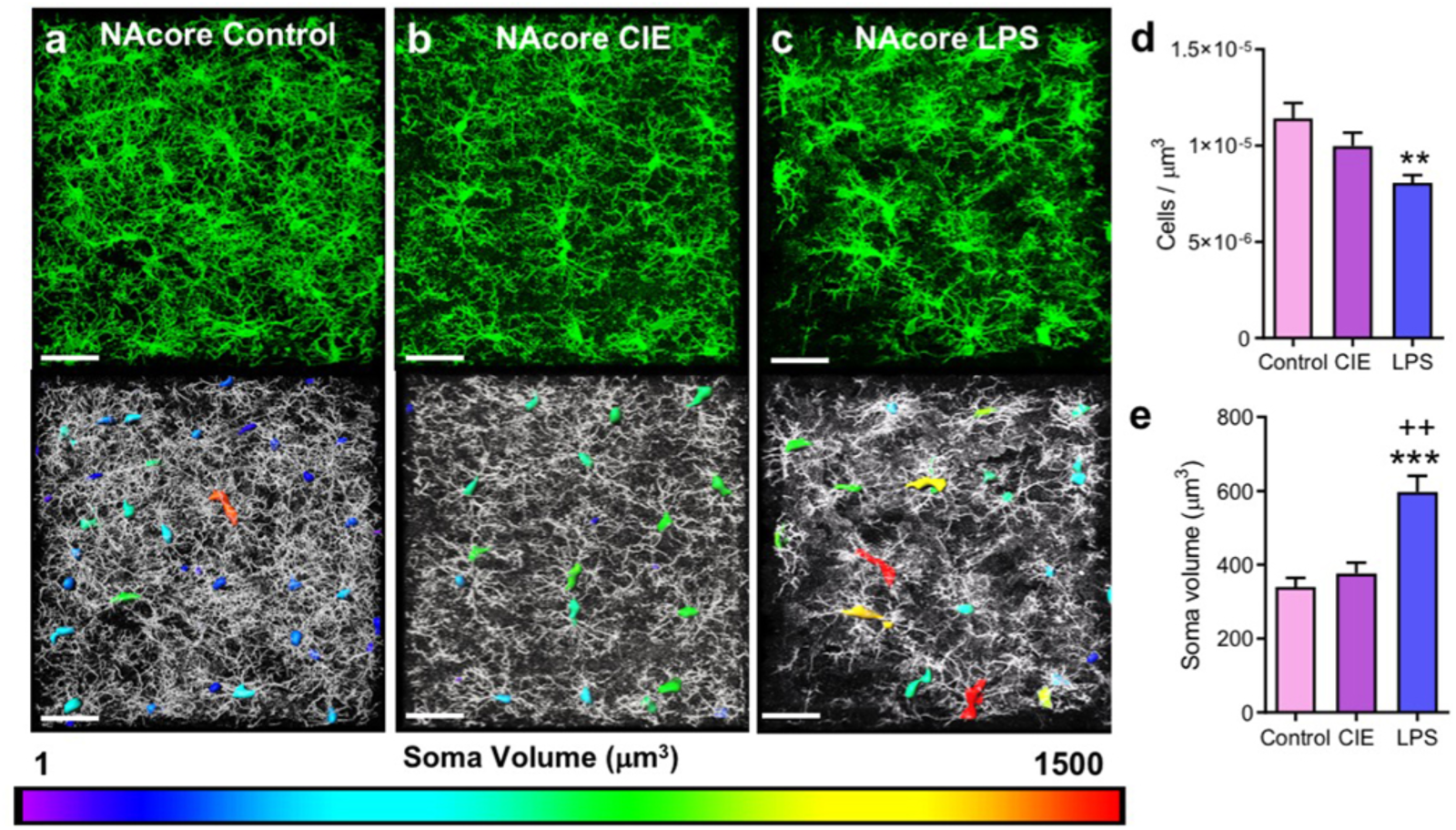
CIE and LPS exposure resulted in an increase in somatic volume of microglia in the NAcore. **a**) Representative Iba-1 staining of microglia in the NAcore of control, (**b**) CIE, and (**c**) LPS-exposed rats. Somas are color-coded to indicate the spectrum of soma volumes observed for each treatment. **d**) LPS-exposed rats exhibited a reduction in the number of Iba-1 immunoreactive microglia per micron3 in the NAcore. **e**) When soma volume was averaged across all images for each animal, only LPS significantly increased the soma volume compared to control rats. \**p*<0.05, \*\*\**p*<0.001 compared to control, ++*p*<0.01 compared to CIE. Scale bar=40 µm.

### Dichotomous changes in microglial morphology and complexity following CIE and LPS exposure in the PL cortex and NAcore revealed by high resolution single cell imaging

To quantify and assess microglial structure reorganization following CIE or LPS exposure, we used high-resolution microscopy with imaging parameters designed to fully capture individual microglia and their corresponding process fields. As shown in the representative images in Figure 6a-c, Iba-1 immunohistochemical detection revealed detailed microglial cells in the PL cortex. Once the imaged cell was digitally isolated and skeletonized in 3D, assessment of the complexity of microglial arbors using Sholl sphere analysis was carried out (radii set at 1 µm intervals from the center of the cell). A two-way repeated measures ANOVA revealed a significant treatment by distance interaction for LPS compared to control rats (F_(49,392)_=3.981, *p*<0.0001) (Figure 6d). Further, Bonferroni-corrected pairwise comparison tests indicated that LPS treated rats had a significantly greater number of Sholl sphere breaks at 11-14 µm from the center of the cell (*p*<0.05). However, there was no significant treatment by distance interaction of the number of intersections for CIE-exposed versus control rats (F_(49,392)_=1.16, *p*=0.216). Isolated individual microglia in the PL were digitally rendered, using space-filling models, to determine microglia cell volume. At the single cell level, space filling models were generated for the entire microglial cell, including both the soma and processes. As is shown in Figure 6e, a one-way ANOVA revealed a significant main effect of treatment on cell volume (F_(2,12)_=22, *p*<0.0001). Tukey’s multiple comparison test further revealed that microglia of CIE-exposed rats had significantly lower overall cellular volume compared to the control group (*p*=0.03). In contrast, LPS-exposed rats exhibited significantly larger overall cellular volume compared to controls (*p*=0.008) and CIE-exposed rats (*p*<0.0001). The intensity of Iba-1 signal within the somatic compartment was quantified and expressed relative to the whole cell intensity. A one-way ANOVA revealed a significant main effect of treatment (F_(2,12)_=28.58, *p*<0.0001, Figure 6f). Tukey’s multiple comparison test indicated that CIE *(p*=0.026) and LPS *(p*<0.0001)-treated rats displayed increased normalized Iba-1 intensity compared to controls, whereas LPS displayed a significantly greater Iba-1 intensity increase when compared to CIE *(p*=0.002).

**Figure 6.**
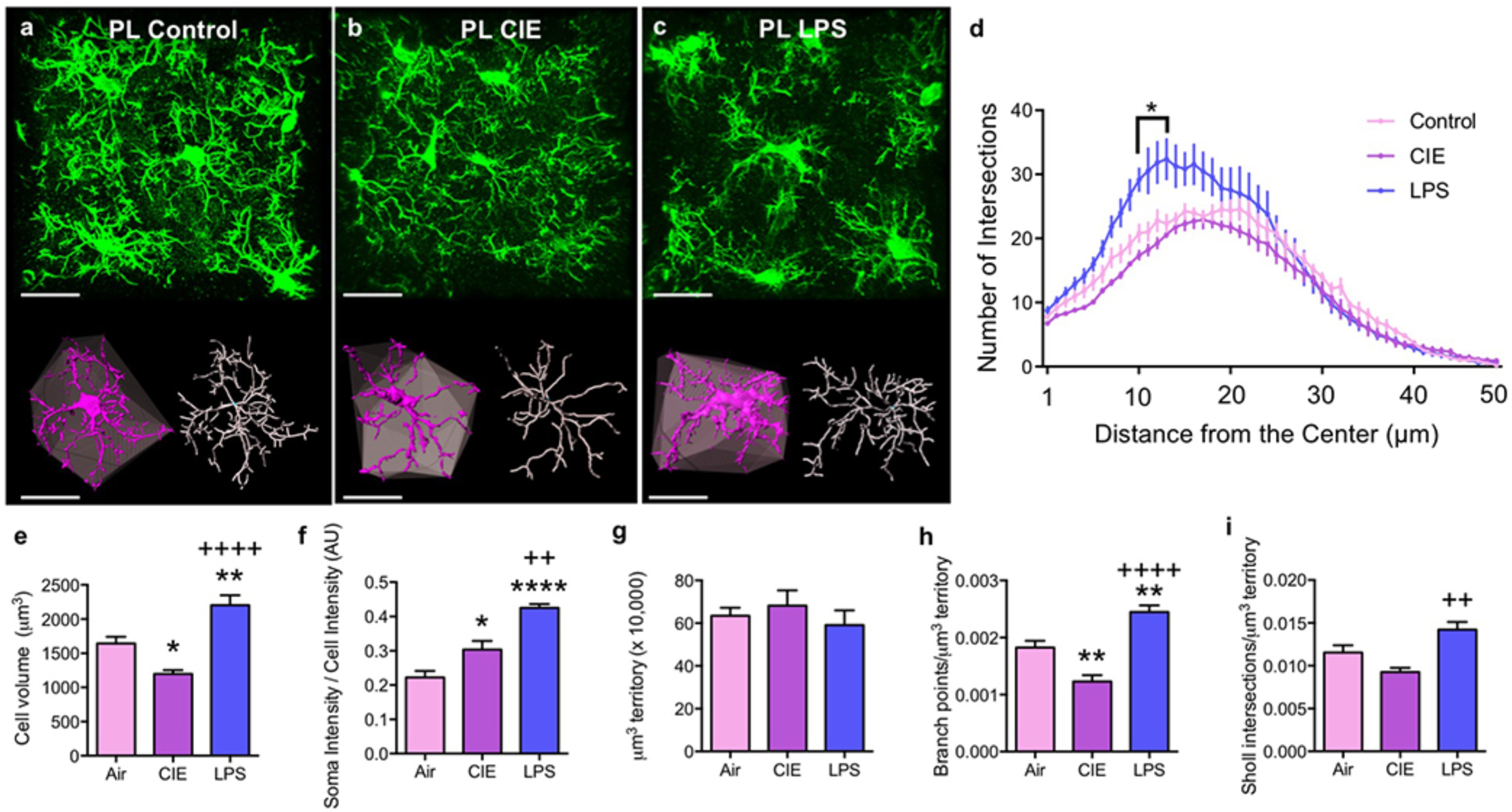
CIE and LPS exposure resulted in dichotomous alterations in microglia complexity in the PL cortex. **a-c**) Raw data (green), digitized single cells (purple) and associated convex hulls (grey), as well as skeletonizations (white) for PL cortex microglia in (**a**) control, (**b**) CIE-exposed, and (**c**) LPS-exposed rats. **d**) Sholl analysis revealed that LPS exposure resulted in a greater number of breaks (in 1 µm intervals) closer to the center of the cell compared to control rats. **e**) There was no difference between groups in the average territory occupied by microglia (‘Hull’ volume). **f**) CIE exposure reduced while LPS exposure increased the average volume of microglia. **g**) When normalized to the hull volume, the reduction in volume following the CIE exposure was lost while the LPS-induced increased in volume was still observed. **h**) There was no effect of CIE or LPS on the number of branch points compared to control rats, yet LPS increased branching when compared to CIE. **i)** When normalized to the convex hull volume, CIE exposure decreased while LPS increased the number of branch points. **j**) When normalized to the hull volume, there was no effect of CIE or LPS in the number of Sholl intersections compared to control rats in the PL cortex, yet LPS was significantly different compared to CIE. \**p*<0.05, \*\**p*<0.01 compared to control, ++*p*<0.01, +++ *p*<0.001, ++++ *p*<0.0001 compared to CIE. Scale bars = 20 µm

Polygonal convex hulls generated from skeletonized microglial processes were then used to assess the spatial territory individual cells occupy. This index has been commonly used as a means to normalize measures of morphological complexity [33]. A one-way ANOVA revealed that there was no difference between groups in the average cellular territory (convex hull) that individual PL cortex microglia occupy within the parenchyma (F_(2,12)_=0.55, *p*=0.589, Figure 6g). Another parameter used to describe microglia structure and complexity is the degree and extent of process branching. As shown in Figure 6h, a one-way ANOVA comparing the number of branch points relative to convex hull volume also revealed a main effect of treatment (F_(2,12)_=29.02, *p*<0.0001). Tukey’s multiple comparison test revealed that CIE exposure reduced branch points (*p*=0.007), whereas LPS increased the number of branch points when compared to both control (*p*=0.006) and CIE-exposed rats (*p*<0.0001). As a final measure of cellular complexity, the total number of Sholl intersections was expressed relative to convex hull volume. A one-way ANOVA revealed a significant main effect of treatment (F_(2,12)_=10.53, *p*=0.002), with Tukey’s multiple comparison test indicating that neither CIE (*p*=0.127) nor LPS (*p*=0.072) exposure significantly decreased the number of Sholl intersections when normalized to the hull volume compared to control rats. However, LPS exposure increased the number of Sholl intersections when compared to CIE rats (*p*=0.002, Figure 6i).

The single cell structural analysis approach was next used to assess morphological properties of microglia in the NAcore. Representative confocal images of the microglia in the NAcore for the three experimental groups are presented in Figure 7a-c. As was observed with microglia in the PL cortex, a two-way repeated measures ANOVA revealed a significant treatment by distance interaction in the number of Sholl intersections in LPS-exposed rats compared to control rats (F_(49,392)_=8.78, *p*<0.0001 Figure 7d). Bonferroni-corrected pairwise comparison tests indicated that LPS exposure significantly increased the number of Sholl intersections at 11-14 and 16-23 microns from the center of the cell (*p*<0.05). Although there was a significant treatment by distance interaction in the number of Sholl intersections in CIE compared to control rats (F_(49,392)_=1.49, *p*=0.022), Bonferroni-corrected pairwise comparison tests indicated there were no significant differences between CIE and control. A one-way ANOVA comparing microglial cell volume between the three groups indicated a significant main effect of treatment (F_(2,12)_=5.21, *p*=0.024, Figure 7e). However, Tukey’s multiple comparison test indicated a significant difference between CIE and LPS-exposed rats (*p*=0.019), but no significant difference for either CIE (*p*=0.341) or LPS (*p*=0.225) compared to controls. As described above, we quantified the intensity of the Iba-1 signal within the somatic compartment and expressed this as relative whole cell intensity. A one-way ANOVA revealed a significant main effect of treatment (F_(2,12)_=8.88, *p*=0.004, Figure 7f). Tukey’s multiple comparison test indicated that LPS increased the normalized Iba-1 intensity compared to control rats (*p*=0.004), yet there was no significant difference between control and CIE-exposed rats (*p*=0.307). Analysis of the convex hull volume in the NAcore revealed that data from control animals was non-normally distributed (W=0.630, *p*=0.001). A Kruskal-Wallis nonparametric one-way ANOVA revealed there were no significant group differences in the extent of territory covered by microglial processes (K-W=3.62, *p*=0.171, Figure 7g). A one-way ANOVA revealed no significant effect of treatment on the number of branch points normalized to convex hull volume (F_(2,12)_=1.55, *p*=0.253, Figure 7h), and the number of Sholl intersections normalized to convex hull volume was also not significantly different between groups (F_(2,12)_=2.77, *p*=0.102, Figure 7i).

**Figure 7.**
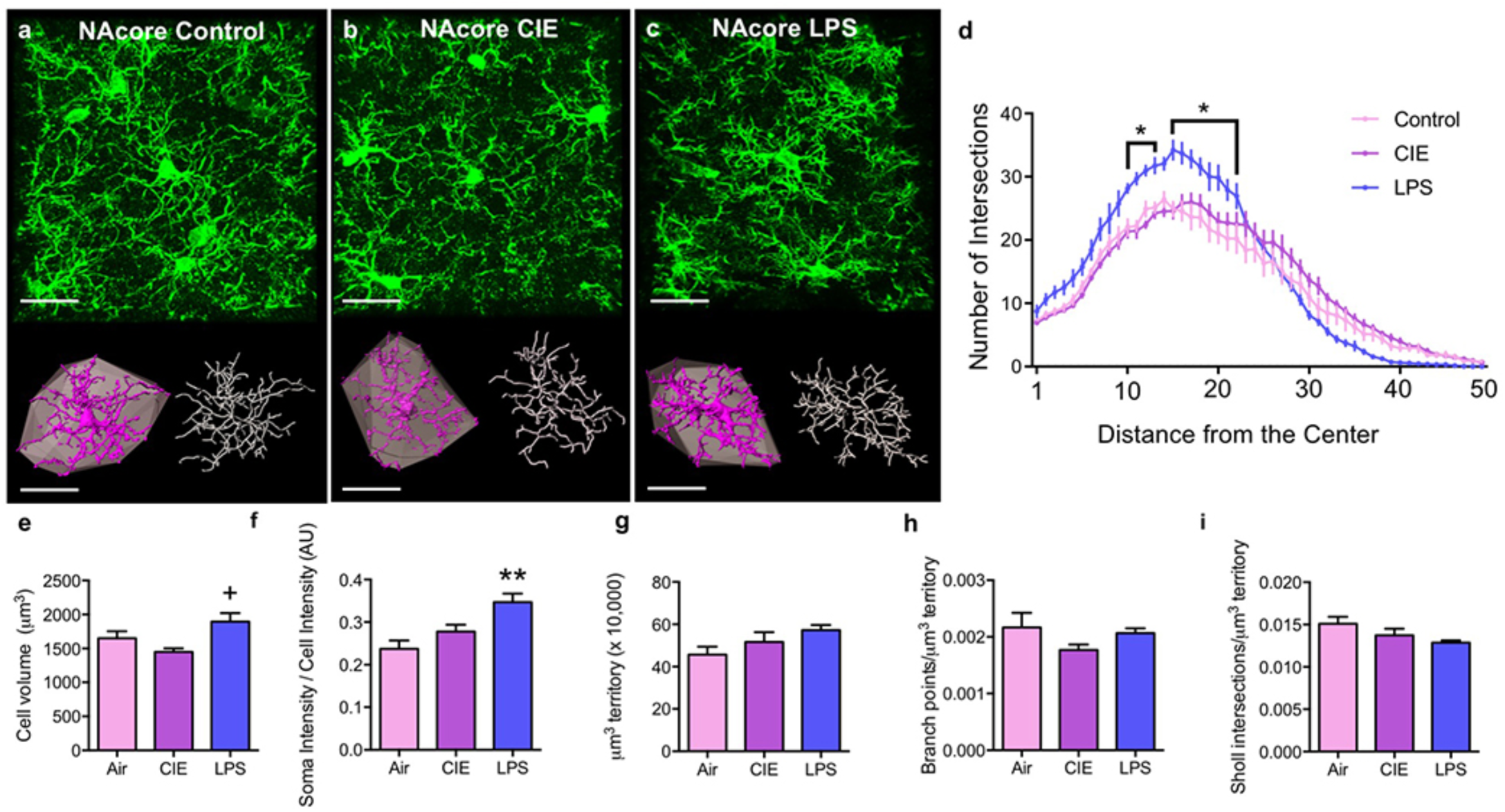
Differential effects of CIE and LPS on microglial complexity were not observed in the NAcore. **a-c**) Raw data (green), digitized single cells (purple) and associated convex hulls (grey), as well as skeletonizations (white) for NAcore microglia in (**a**) control (**b**) CIE-exposed and (**c**) LPS-exposed rats. **d**) Sholl analysis revealed that LPS-exposure resulted in a greater number of breaks (in 1 µm intervals) closer to the center of the cell compared to control rats. **e**) There was no difference between groups in the microglia convex hull volume in the NAcore. **f**) Neither CIE nor LPS exposure affected microglial volume in the NAcore compared to control. There was no difference between groups in microglia cell volume normalized to the volume of territory occupied (**g**) or **h**) the number of branch points. **i**) Uniquely, in the NAcore LPS exposure significantly decreased the number of branch points normalized to the volume of territory occupied when compared to control rats. **j**) The number of Sholl intersections normalized to the hull volume was not different between groups in the NAcore. \**p*<0.05 compared to control, +*p*<0.05 compared to CIE. Scale bars = 20 µm

## Discussion

The results of the current study revealed that CIE and LPS exposure differentially alter the morphological profile of microglia, as revealed by immunohistochemical detection of Iba-1, 3D digital reconstruction of confocal images, and analysis of morphometric properties. Rats that were rendered dependent on ethanol by 15 days of CIE exposure and sacrificed 10 hours into withdrawal exhibited brain region-dependent alterations in morphologic properties of microglia that were distinct from those induced by LPS in both the PL and NAcore, two relevant nodes of the addiction and cognitive neurocircuitry [52, 55]. When fields of microglia in the PL were sampled in rats exposed to LPS, we observed an increase in microglial somatic volume compared to controls and CIE-exposed rats. Consistent with what others have shown for alcohol [56] and LPS [57] exposure, high magnification confocal imaging of individual microglia demonstrated that both CIE and LPS exposure increased Iba-1 intensity within the somatic compartment, an effect consistent with microglial activation [58]. Interestingly, CIE-, but not LPS-mediated increases in somatic Iba-1 intensity were specific to the PL cortex. LPS exposure significantly increased various indices of PL microglial complexity (overall cell volume, branch points/um3 territory), yet PL microglia from CIE-exposed rats exhibited reductions in these same indices. Further, we observed that morphological alterations in the NAcore due to both LPS and CIE exposure were of substantially lower magnitude when compared to the PL, indicating the observed effects are brain region dependent. This is likely not due to basal differences in the ability of Iba-1 to fully label microglia in these two brain regions given that comparisons of Cx3cr1-EYFP and Iba-1 labeling in both the PL cortex and NAcore produced nearly identical results, with Iba-1 actually producing slightly more complete labeling. Taken together, these data suggest that CIE and LPS produce dichotomous alterations in microglial process architecture in which CIE simplified and LPS increased microglia process branching. These distinct morphometric signatures may reflect unique aspects of the correspondingly distinct pathophysiologies engaged by CIE and withdrawal, relative to LPS exposure. Moreover, these findings suggest that careful analyses of microglia architecture using a single cell imaging approach can reveal discrete alterations in microglia morphological complexity that could be overlooked with more coarse analyses of parameters like soma size and Iba-1 intensity.

### LPS has heterogenous effects on somatic volume and complexity

LPS exposure has previously been shown to activate microglia by binding to TLR4 and engaging the innate immune response, ultimately driving increased cytokine release. However, the degree to which LPS engages alterations in microglia morphometric properties is highly dependent on the dose and timeframe. Consistent with our data, LPS exposure at doses similar to what was used in the present study produced a hyper-ramified morphological profile characterized by an increased soma size and increased branching [33, 59]. Activated microglia that display the hyper-ramified structural profile have also been observed following chronic stress [60], ischemic stroke [41], and traumatic brain injury [33]. In addition, we show that LPS-induced increases in soma size occurring in the PL cortex are also associated with increased overall cell volume in conjunction with the increased branching complexity and number of branch points.

Our data are largely consistent with a previous study showing that 24 hours after exposure to a low dose LPS microglia soma volume is increased in the mPFC with no alteration in the number of cells [61]. Also consistent with our results, this study demonstrated that LPS increased process thickness and branching complexity [61]. However, our data provide new evidence that LPS-mediated increases in soma volume are brain region-independent, whereas increases in branching patterns were largely brain region-dependent given that they were not observed in the NAcore. Taken together, these findings indicate that enlarged soma volume and enhanced Iba-1 intensity coincide with heightened microglia activation. However, the brain region-specificity revealed by our data suggest that branching patterns are likely a more nuanced metric that may reflect unique aspects of a specific activation profile. Taken together, our data indicate that LPS has a greater impact on PL cortical microglia morphology compared to microglia in the NAcore. Given that the PL cortex regulates multiple cognitive processes [62], and that frontal cortical-dependent processing of emotionally-relevant stimuli is disrupted following LPS administration in humans [63], this finding suggests that the PL cortex-specific alterations in microglial complexity following LPS may play a role in the cognitive decline following exposure to environmental insults such as TBI.

### CIE-induced alterations in microglia branching complexity occur in a brain region-dependent manner

At a global level, the frontal cortex and NAc exhibit anatomically unique gene expression profiles in humans with alcohol use disorder [64], including genes linked to the induction of neuroinflammatory responses [65]. Specifically, within the subset of alcohol-responsive genes, only 6% are shared between the frontal cortex and NAc [66]. Consistent with this human data, CIE in rodents alters the expression level of a larger amount of alcohol-responsive genes in the PL cortex when compared to the NAcore [67]. The data presented here are also consistent with the hypothesis that the microglia in the PL cortex are more responsive to CIE when compared to microglia in the NAcore. We demonstrate that simplifications in microglia process density during withdrawal from CIE, a microglial morphological signature commonly observed in aged rodents [68], occur in the PL cortex and not the NAcore. Thus, the greater impact of alcohol exposure on gene expression in the cortex may contribute to the brain region specificity of the effects on microglial cellular architecture described in our study. However, whether these brain region-specific morphological differences translate to functionally differential characteristics of microglial activation remains to be fully determined. Given that microglia play an important role in shaping synaptic plasticity by releasing neuroimmune modulators [69], our data are consistent with the suggestion that microglial-mediated neural circuit adaptation following chronic ethanol exposure and withdrawal could be mechanistically distinct in the PL cortex when compared to the NAcore.

### 4.3 Comparing withdrawal from CIE- and LPS-mediated microglial activation

Microglial activation has been implicated in the regulation of the neural processes linked to cognition, memory, and associative learning, as well as with the neuroplasticity and neuropathology associated with acute and chronic exposure to alcohol [25, 46, 70–73]. While it is clear that LPS-induced activation of microglia via TLR4 leads to an induction of the proinflammatory cytokine TNFα, the ability of ethanol exposure to do the same is currently under debate [23, 25]. As an example, prolonged (6 months) of ethanol exposure was observed to increase the overall proportion of activated microglia relative to total microglia in the hippocampus, yet did not impact TNFα expression [22]. Furthermore, binge-like alcohol exposure during adolescence increased CD11b immunoreactivity in the hippocampus, an additional measure of microglial activation [74], yet did not alter TNFα expression [11]. This differential impact of ethanol and LPS on microglia TNFα expression may explain dichotomous impacts of ethanol exposure and LPS exposure on overall microglia volume or branching patterns.

An emerging concept that is gaining support is that ethanol exposure produces a state of “partial activation” of microglia [11, 26]. Given that LPS increased both somatic Iba-1 intensity and soma volume to a greater extent than CIE, our data support this hypothesis. However, at the single cell level, we observed that microglia display two disparate and unique morphological signatures following CIE and withdrawal or LPS exposure. While we observed a decreased number of microglia, reduced overall cell volume and decreased branching following CIE there was an increase in overall cell volume and process density following exposure to LPS but no alteration in the total number of microglia. Thus, an alternative hypothesis is that CIE leads to hypoactivation of microglia in the PL cortex. In our data set, reduced process density and overall cell volume could serve as evidence for hypoactivation. This interpretation would suggest that following chronic alcohol exposure, microglia in the PL cortex may be less responsive to activation and thus less suited to fulfill their protective role following additional environmental insults [75]. This may be reflected as reduced ability to release cytokines, decreased overall microglial process mobility, and disrupted phagocytosis [76].

As described above, we observed reduced microglia cell volume and branching following CIE exposure in the PL cortex. This represents a structural profile similar to what has been observed in the neocortex of Alzheimers patients [77] and in rodent models of aging [68, 78]. In contrast, after two 1mg/kg systemic administrations of LPS separated by 24 hours, we observed increased microglia cell volume and branching. These dichotomous morphological signatures could also be consistent with another unique hypothesis, whereby LPS- and CIE-mediated microglial activation produce mechanistically distinct gene expression and cytokine and chemokine release patterns. Consistent with this hypothesis, binge alcohol exposure during adolescence leads to increased microglia proliferation during adulthood, independent of phagocytic activity or increased MHC-II expression commonly associated with ‘activated’ microglia [79]. Thus, ethanol-induced microglia activation is likely independent of the innate immune reaction observed in microglia following LPS treatment (i.e. upregulation of MHC-II) [80]. Additional experimentation is required to determine if CIE and withdrawal produces partial activation, hypoactivation, or a mechanistically unique activation state in PL cortical microglia. However, in light of the ongoing revisions to the cellular, molecular, and structural characteristics associated with microglia activation states [81], the use of high resolution microscopy and digital reconstruction will no doubt provide new structural biomarkers that can be used to better understand how alterations in cellular complexity relate to microglial function.

## Conclusion

LPS had a profound impact on soma volume in both the PL and NAcore when compared to CIE. However, CIE exposure reduced multiple indices of microglia complexity whereas LPS increased such measurements; an effect that was relegated to the PL cortex. Interestingly, our examination of microglial morphological profiles at high magnification revealed that CIE exposure produced a structural profile that is similar to what has been reported in Alzheimer’s disease [77] and in rodent models of aging [68, 78]. In contrast, LPS produced a structural profile that is more similar to what has been observed following chronic stress [60], ischemic stroke [41], and traumatic brain injury [33]. The present study may lay a framework for high fidelity imaging and 3D structural characterization following distinct environmental insults, and that usage of the methods described herein will allow for a better understanding of the scope of microglial diversity through high structural characterization and classification

## Acknowledgements

This work was supported by R00 DA040004 (MDS), R01 AA010983 (LJC), R01 AA022701 (LJC), and T32 DA007288 (BMS). The authors thank Dr. Christopher Cowan and Catherine Bridges for providing *Cx3cr1*^*CreER*^-EYFP transgenic mice.

## Author contributions

BMS, JDL, JAM, and KNH conducted study investigation. BMS, JDL, LJC, and MDS participated in writing, reviewing, and editing. BMS, LJC, and MDS provided conceptualization for all experiments. BMS and MDS designed the methodology for all analyses. LJC and MDS provided supervision over the conducted research. BMS and MDS participated in visualization of all data.

---

**Figure S1.**
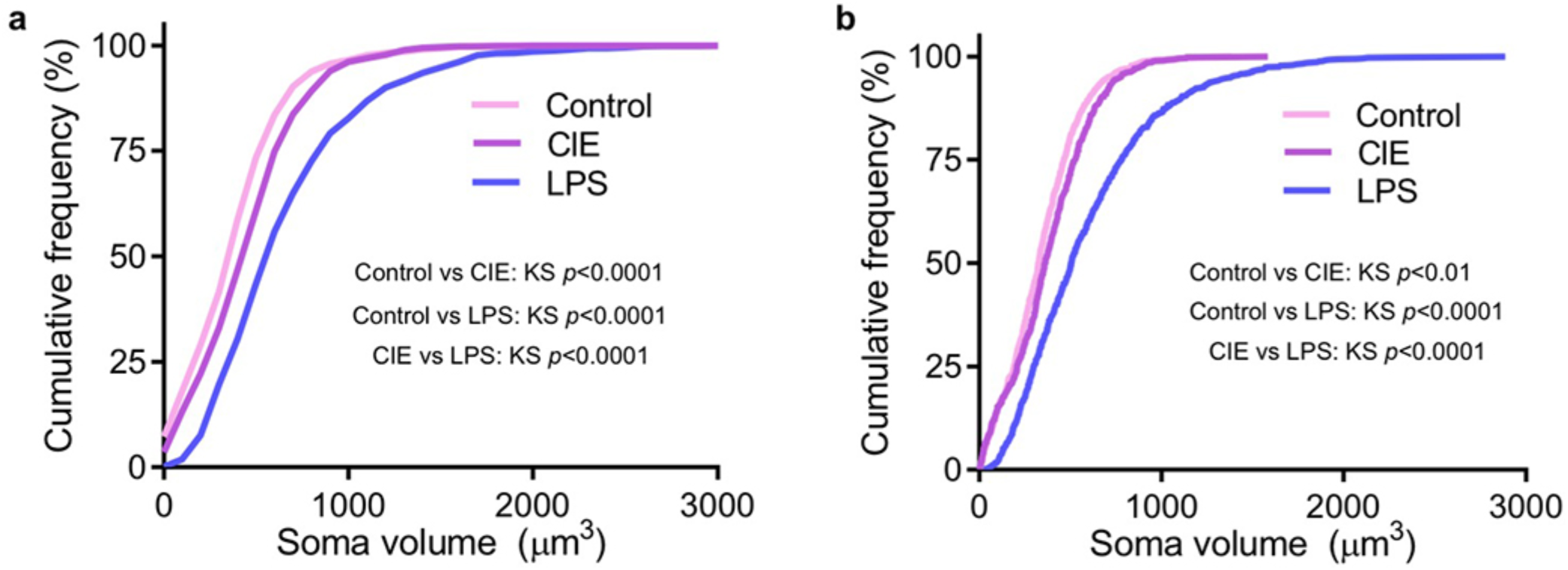
Cumulative frequency distributions for Control, CIE-, and LPS-exposed rats in PL and NAcore. **a-b)** Examination of the frequency distribution of somatic volume revealed that both CIE and LPS exposure resulted in a rightward shift in soma volume distribution curve in the PL **(a)** and NAcore **(b)**.

